# PairK: Pairwise k-mer alignment for quantifying protein motif conservation in disordered regions

**DOI:** 10.1101/2024.07.23.604860

**Authors:** Jackson C. Halpin, Amy E. Keating

## Abstract

Protein-protein interactions are often mediated by a modular peptide recognition domain binding to a short linear motif (SLiM) in the disordered region of another protein. The ability to predict domain-SLiM interactions would allow researchers to map protein interaction networks, predict the effects of perturbations to those networks, and develop biologically meaningful hypotheses. Unfortunately, sequence database searches for SLiMs generally yield mostly biologically irrelevant motif matches or false positives. To improve the prediction of novel SLiM interactions, researchers employ filters to discriminate between biologically relevant and improbable motif matches. One promising criterion for identifying biologically relevant SLiMs is the sequence conservation of the motif, exploiting the fact that functional motifs are more likely to be conserved than spurious motif matches. However, the difficulty of aligning disordered regions has significantly hampered the utility of this approach. We present PairK (pairwise k-mer alignment), an MSA-free method to quantify motif conservation in disordered regions. PairK outperforms both standard MSA-based conservation scores and a modern LLM-based conservation score predictor on the task of identifying biologically important motif instances. PairK can quantify conservation over wider phylogenetic distances than MSAs, indicating that SLiMs may be more conserved than is implied by MSA-based metrics. PairK is available as open-source code at https://github.com/jacksonh1/pairk.

## INTRODUCTION

Protein-protein interactions often involve a modular peptide recognition domain in one protein binding to a short linear motif (SLiM) located in the disordered region of another protein. Examples of such peptide recognition domains, or SLiM-binding domains, include SH3, SH2, and PDZ domains. Over several decades, researchers have defined core motifs for many SLiM-binding domains that describe the minimal sequence requirements for a peptide to bind to the domain (*1*). These typically span 3-10 residues, with only a few positions fully defined, and are often described using regular expressions that list the residue(s) required or allowed at each position. The low sequence complexity of these motifs does not typically provide enough information to map specific biological interactions using sequence alone. For example, when searching the proteome for sequences that match a motif regular expression, thousands of instances are usually found. However, the vast majority of these matches are not biologically relevant, making the prediction of new interactions challenging.

One reason that SLiM definitions fail to predict biologically relevant interactions is that the core binding motif alone is often insufficient for binding. Interactions between SLiMs and their cognate SLiM-binding domains display diverse binding mechanisms that extend beyond the core motif to enhance binding or confer specificity. For instance, some SLiM interactions use sequence flanking the core motif (*2, 3*), a secondary binding site on the binding domain (*4*), or tandem motif repeats to increase affinity via avidity (*5–7*). Moreover, different binding partners can use distinct mechanisms to recruit the same SLiM-binding domain (*2, 3, 8, 9*), making it impossible to define a single motif that captures all the information required for binding. Consequently, inferring the biological relevance of any potential SLiM from the core sequence motif is difficult.

In the bioinformatic analysis of candidate SLiM interactions, filters or annotations are often employed to discriminate between biologically relevant and improbable motif matches (*10*). Predicted disorder, e.g. using IUPRED scores (*11, 12*), is commonly used to rule out matches within globular domains, and functional annotations from Gene Ontology (*13, 14*) can identify candidate motifs that share functions with a binding domain of interest (*10*).

A promising criterion for identifying biologically pertinent SLiMs is the evolutionary conservation of the motif. Filtering based on conservation presumes that functional motifs are more likely to be conserved than are spurious motif matches. Despite SLiMs having the potential for rapid evolution (*15*), genuine motifs are often more conserved than their surrounding sequences and frequently emerge as islands of conservation amidst highly variable disordered regions (*16–19*). Methods such as SLiMPrints (*16*) have been devised to quantify relative local motif conservation. Assigning a conservation score to a SLiM instance typically involves collecting sequences homologous to the protein containing the SLiM, creating a multiple sequence alignment (MSA), and then quantifying how conserved potential motif residues are compared to adjacent residues (*16, 19*).

A recognized limitation of current SLiM conservation metrics is their reliance on an initial MSA (*16, 18*). Compared to folded domains, intrinsically disordered regions (IDRs) are often extremely challenging, if not impossible, to align. MSAs are constructed under the assumption that residues in each column of the alignment correspond to directly comparable structural positions across sequences. Folded domains, due to strict evolutionary constraints on structure, often exhibit this type of conserved residue positioning across homologs of the same domain. Consequently, MSAs of folded domains are highly informative. In contrast, positions across homologous IDRs are often not comparable because IDRs evolve differently than folded domains. Homologous IDRs often exhibit characteristics such as unique residue composition, a high prevalence of insertions and deletions, and low-complexity or repetitive regions that make alignment challenging (*20, 21*). Resulting MSAs are often difficult to interpret and can be misleading.

Progress has been made in developing alignment-free methods for comparing homologous disordered regions. Although not specifically designed for IDRs, k-mer based sequence distance methods can calculate the similarity between two IDRs without an alignment. These methods work by comparing the counts or frequencies of unique subsequences of length *k* within each sequence (*22*). A variation of a k-mer method, SHARK was recently developed specifically for identifying homologous IDRs (*23*). It utilizes k-mer physicochemical similarity to identify homologous IDRs with better performance than standard methods. Though useful for quantifying IDR similarity, k-mer based distance methods cannot be used for residue-level conservation scores directly. Recent advances in protein large language models (LLMs) have led to methods for predicting conservation scores without the need for a input MSA or set of homologous sequences (*24, 25*). Yeung et al. found that their conservation score predictor, Kibby, predicted higher conservation for known phosphorylation sites in disordered regions than traditional conservation scores from MSAs (*24*).

Despite the lack of conservation in the positioning of residues in IDRs, other sequence features can be conserved. These include the patterning and abundance of specific residues, and the distribution of physicochemical properties across the sequence (*26–32*). Several alignment-free methods have been developed to quantify bulk properties of IDRs and their conservation (*28, 33, 34*). In contrast to bulk physicochemical properties, which can be conserved without conserving the precise positioning of residues, functional short linear motifs (SLiMs) must form specific contacts with residues in the binding pockets of their cognate SLiM-binding domains. Therefore, SLiMs are distinct from other functional properties of IDRs. Currently, there is a lack of alignment-free tools developed to quantify the conservation of SLiMs in disordered regions.

We developed a simple, MSA-free method—PairK (pairwise k-mer alignment)—for quantifying residue-level conservation of short motifs in IDRs. PairK can be applied to raw sequences or to sequences embedded using ESM2 (*35*). We developed a benchmark to evaluate the effectiveness of our method by using it to distinguish experimentally verified SLiMs vs. background motif matches in human disordered sequences. Our method outperformed both standard MSA-based and modern LLM-based conservation score predictors like the Kibby method of Yeung et al. (*24*). We found that PairK can quantify conservation across broader phylogenetic distances than MSAs and are more effective at distinguishing verified SLiMs from background matches over these distances. This suggests SLiMs may be more conserved—and evolve more slowly—than previously believed. PairK is available as an open source package, free for use by all, at – https://github.com/jacksonh1/pairk.

## RESULTS

### Sequence alignments of disordered regions confound the quantification of SLiM conservation

To evaluate different methods for quantifying conservation, we first developed a pipeline to retrieve and process homologous sequences for a specified protein. We used OrthoDB (*36*) to obtain precompiled groups of homologous sequences at various phylogenetic levels (called orthologous groups in OrthoDB). For a protein of interest (the query protein), the pipeline performs the following steps: it locates the query protein in OrthoDB, retrieves the orthologous group at a specified phylogenetic level, compares each homolog to the query protein and selects the least divergent homolog in each organism, clusters the least divergent homologs to reduce redundancy, and finally creates an MSA from the remaining sequences (see Methods for details).

To explore potential issues arising from aligning disordered regions, we examined the MSAs of proteins with verified SLiMs. Figure 1A presents a section of an MSA of RIAM, a verified Ena/VASP EVH1 binding partner in humans (*37*), aligned with its vertebrate homologs. The folded domain exhibits high conservation and minimal gaps, while the disordered region containing the Ena/VASP EVH1 binding motif (LPPPP) displays a larger number of insertions/deletions and a higher overall apparent divergence. Figure 1B provides a detailed view of the MSA region containing the SLiM. Although all the homolog sequences have at least one motif match in the short region shown, few are well-aligned with the human RIAM motif, resulting in an artificially low conservation score for the first motif position. Several positions in the alignment, including several in the SLiM, show artificially high conservation due to prolines that are not confidently aligned (indicated by red arrows). This is because prolines are assigned a high score in most substitution matrices, and disordered regions are rich in proline residues. To optimize the alignment score, alignment algorithms tend to align prolines in IDRs when there is no other strong signal.

**Figure 1.**
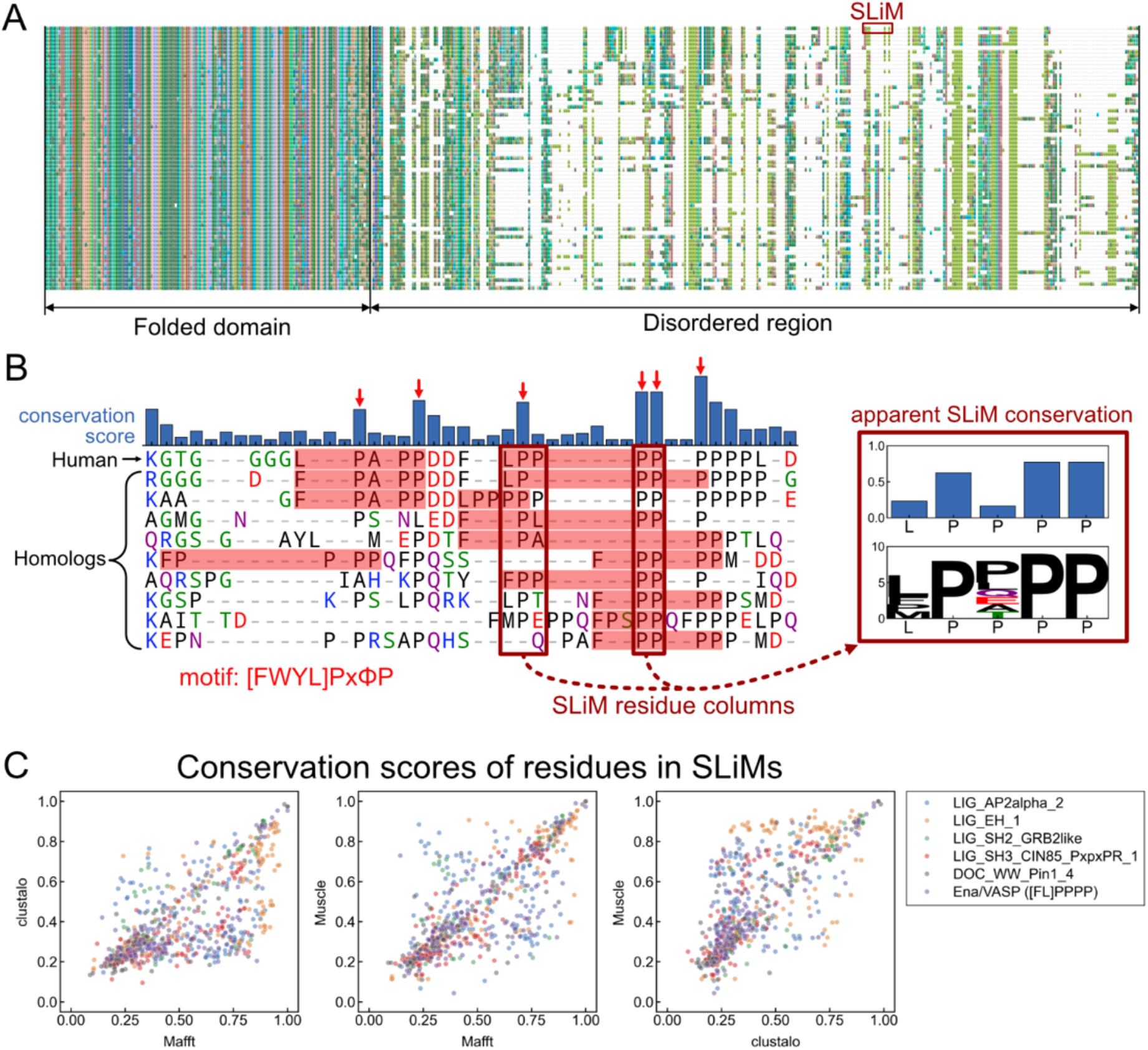
Detecting evolutionary conservation of short linear motifs is confounded by poor alignment of disordered regions. (A) Slice of an MSA of RIAM, which contains an LPPPP Ena/VASP binding motif (*37*), aligned to its vertebrate homolog sequences. (B) *Left* – Part of the SLiM region of the MSA from (A). Matches to the EVH1 motif are highlighted in red. Columns that appear artificially conserved are indicated with a red arrow. *Right* - The apparent conservation of the SLiM residues extracted from the example MSA. X-axis labels are residues in the human protein. White space in the sequence logo indicates gaps in the corresponding alignment columns. Bar plot shows the conservation scores of the aligned columns. (C) The conservation scores (Shannon entropy from Capra et. al (*42*)) of residues in experimentally verified SLiMs vary with alignment algorithms. Data are from 236 verified SLiM instances (721 residues). Homologs are from metazoans.

To assess the sensitivity of the apparent evolutionary conservation of SLiMs to the underlying MSA, we collected a set of experimentally verified SLiM instances from the Eukaryotic Linear Motif (ELM) database (*38*) or manually curated them from the literature (see Methods), and processed them through the pipeline described. For each set of homologs, we generated MSAs using Mafft (*39*), Muscle (*40*), and Clustalo (*41*) and calculated conservation scores as described in the methods for residues within the verified motifs. Figure 1C demonstrates that the conservation scores of many residues vary significantly when using different alignments, suggesting that the MSA, and thus the resulting motif conservation score, is poorly determined.

### PairK (pairwise k-mer alignment)

As an alternative to using MSAs for quantifying the conservation of SLiMs, we developed a model that assumes homologous IDRs are globally unconserved but contain short, positionally conserved stretches of residues (SLiMs). Based on this assumption, we developed a scoring method to quantify the relative conservation of short sequences in IDRs, which we term PairK (pairwise k-mer alignment) (illustrated in Figure 2A).

**Figure 2.**
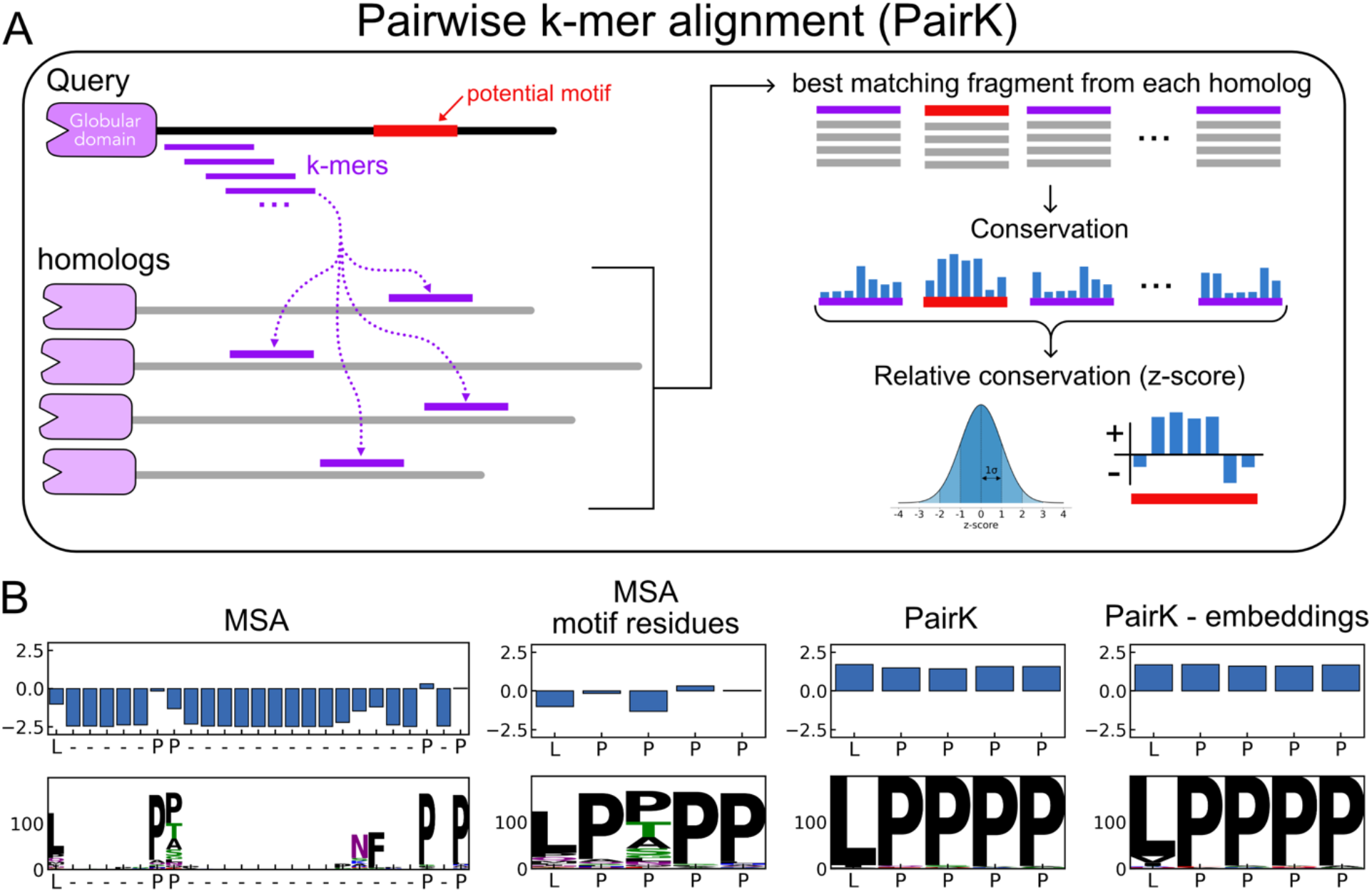
The pairwise k-mer alignment method (PairK) for quantifying the conservation of SLiMs. (A) Schematic of the method. (B) Example z-scores and sequence logos for an Ena/VASP binding motif from the protein RIAM and its vertebrate homologs. X-axis labels are residues in the human protein. White space in the sequence logos indicates gaps in the corresponding alignment. The results using an MSA (the same MSA from Figure 1A) are shown at *left*. The positions in the MSA corresponding to the SLiM residues in the human sequence (LPPPP) are extracted and shown in the *middle-left* panel, with gaps removed. The results from PairK (*middle*-*right*) and the embedding-based variant of PairK (*right*) suggest that the LPPPP motif is more conserved than it appears in the MSA.

The scoring procedure works as follows (see methods for details). For a query protein with a disordered region containing a potential SLiM, we retrieve homologous sequences using OrthoDB and the pipeline described in the previous section. The IDR of the query sequence is then divided into all possible k-mers, where each k-mer is a subsequence of the query IDR of length k. Each k-mer is then aligned to each homologous IDR in a pairwise manner without gaps, and the best matching subsequence from each homologous IDR, as determined by a scoring matrix, is retrieved. The list of matches, with each subsequence aligned to the query, constitutes a pseudo MSA for each k-mer. We use the term “pseudo MSA” because there is no attempt to perform a global alignment of the retrieved k-mers. Conservation scores are then calculated column-wise for each position in each k-mer pseudo MSA.

To account for the divergence of the motif-containing IDR, we compared k-mer conservation scores for a putative motif to the conservation of all other residues in all k-mer pseudo MSAs for the same IDR, using a z-score. To directly compare PairK scores with MSA scores, we also converted MSA conservation scores to z-scores, such that each final score is relative to the other columns in the IDR region of the MSA. This z-score normalization is very similar to that used in the SLiMprints method (*16*), except that we normalized relative to the entire IDR instead of a window around each residue.

Figure 2B displays the resulting sequence logos and conservation scores for the example SLiM shown in Figure 1, using both the MSA method and PairK. In these logos, the height of each letter indicates the count of that residue in the corresponding alignment column. Gaps in an alignment column result in whitespace above the letters (where the logo letters do not fill the column). For the example in Figure 2B, the motif residues receive much higher scores using the pairwise k-mer method compared to the MSA. Additionally, the sequence logos for the pairwise k-mer method suggest that the L in the first position and the P in the third position are more conserved than is reflected in the MSA analysis.

We developed a variation on PairK that incorporates residue embeddings from ESM2, a large language model trained on protein sequences (*35*). ESM2 residue embeddings are computed for the full-length query sequences and their homologs. When the query IDR is split into k-mers, the corresponding residue embeddings are also sliced out of the query sequence embedding tensor. For each k-mer, the best matching subsequence in a homologous IDR is determined by calculating the Euclidean distance between the associated k-mer embedding slice and the embedding slices of all subsequences of length *k* in the homologous IDR. The subsequence with the lowest embedding distance is selected as the best-matching sequence and used to construct the pseudo MSA. See Figure 2B for an example of applying this approach.

### PairK outperforms other methods for SLiM conservation quantification

Given initially promising results using PairK, as seen in Figure 2B, we sought a systematic approach to compare different methods. We created a benchmark to evaluate the performance of various scoring methods in quantifying SLiM conservation (Figure 3A). We handpicked seven SLiMs based on their abundant experimentally verified instances annotated in ELM (those defined for binding to domains AP2, EH, SH2, SH3, WW, 14-3-3, and Ena/VASP EVH1) (*38*). Using the regular expressions that define these motifs, sourced from the ELM database or the literature, we identified motif matches in disordered regions of human proteins. The matches were divided into two categories: high-confidence verified SLiMs (true positives) and background motif matches. The high confidence verified SLiMs are annotated true positives from the ELM or were manually curated from the literature (Table S2 and S3, see Methods). True positives in the ELM are SLiM interactions that are manually curated and supported by various lines of evidence, primarily experimental (*38*). As in other studies with similar benchmarks (*16, 18*), we assumed that the large majority of background proteome motif matches are not biologically relevant binders, and the fraction of real motifs is much larger in the true positive set than the background set. Our benchmark is based on the expectation that true positives are more evolutionarily conserved, on average, than background motifs.

**Figure 3.**
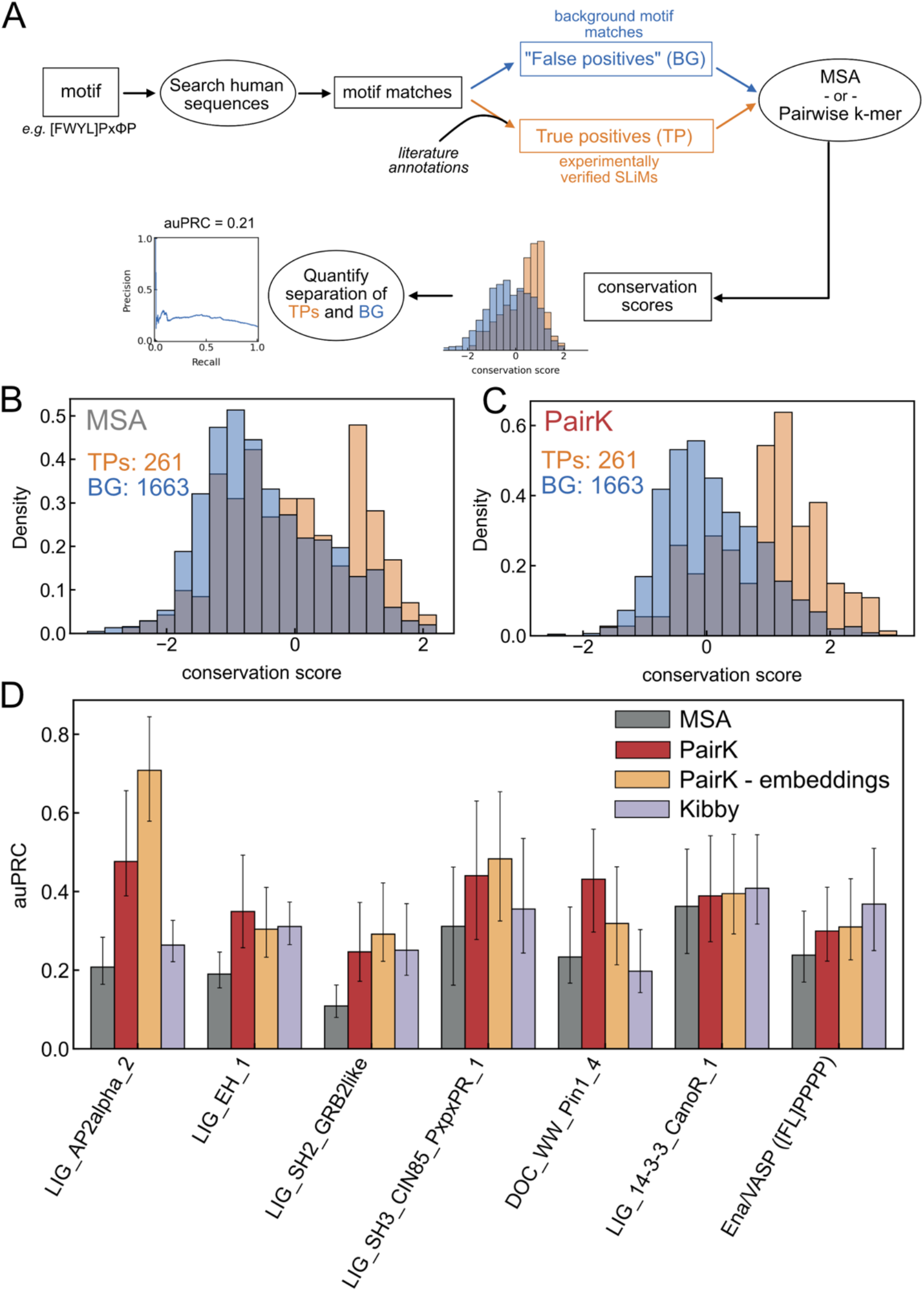
SLiM conservation scoring benchmark. (A) Schematic of the benchmark pipeline. Distributions of the conservation scores of the motifs in the benchmark are shown for the MSA method (B) and PairK (C). Homologous sequences were gathered at the metazoan level. PairK better separates motif matches that are validated (TPs, orange) from background motif matches in the proteome (BG, blue). (D) For each motif in the benchmark, PairK (red) performs better than the MSA method (gray). Error bars are 95% confidence intervals from a bootstrap analysis. The plot is for homologs at the Metazoa level.

Each human protein motif was evaluated using different scoring methods to calculate a conservation score for the query motif, defined as the average conservation score for defined motif residues. We then analyzed how effectively conservation scores distinguish true positives from the background matches, using the area under the precision-recall curve (auPRC) as our performance metric (*43*). For further details, see Methods and Supplementary Table S1.

For each SLiM class in the benchmark, PairK outperforms the MSA method (Figure 3B-D). Interestingly, the degree of improvement varies among the SLiMs. AP2 shows the most significant enhancement (∼2x increase in auPRC), while 14-3-3 shows the least. The embedded version of PairK further improved performance for some SLiMs, with AP2 showing a ∼3x increase in auPRC compared to MSA. For our initial analysis, we set *k* equal to the length of the motif. We also explored the impact of including flanking sequence around the motif match, *i*.*e*., we increased the value of k. In these cases, we performed the pairwise alignment step with the increased value of *k* and then extracted just the core motif residue scores (Figure S1A). Although adding flanking sequence slightly reduced performance, results were still better than for the MSA method for most SLiMs (Figure S1B). The choice of scoring matrix for the pairwise alignment step, including a matrix developed for alignment of disordered regions (*44*), had negligible effects on performance (Figure S1C).

We assessed conservation as calculated using the alignment-free score predictor of Yeung et al., referred to as the Kibby method here. After converting the Kibby scores to z-scores, as for MSA scores, we found that Kibby outperformed MSAs in most instances except for the WW motif (Figure 3D). PairK outperformed Kibby for most SLiMs but was less effective for 14-3-3 and Ena/VASP EVH1.

### PairK quantifies SLiM conservation at greater phylogenetic distances

We examined how various conservation methods performed for orthologous groups at different phylogenetic levels. For progressively more divergent phylogenetic levels (Tetrapoda, Vertebrata, Metazoa) (Figure 4A), the performance of the MSA method remained approximately the same, whereas that of PairK increased (Figure 4B-C, Figure S3). The increasing performance of PairK with increasing phylogenetic level is consistent with greater conservation of the SLiM relative to the rest of the IDR. Figure 4D shows an example. Here we added flanking residues around the motif prior to the pairwise k-mer alignment (increased *k* to 15) to show conservation of the surrounding sequence. The sequence logos and scores look similar by both the MSA and PairK methods at the Vertebrata level. However, the MSA at the Metazoa level suggests that the SLiM is not conserved in this group of species. The MSA-based logo for Metazoa has a large fraction of gaps and includes threonine at the first position. The apparent higher conservation of the second, third, and fourth prolines is likely due to spuriously aligned prolines from poorly aligned sequences. Overall, the MSA suggests that the motif is not conserved in the >200 additional organisms present in the metazoan orthologous group. In contrast, by PairK, the SLiM residues score even higher at the Metazoa level than the Vertebrata level. The sequence logo suggests that the motif residues remain generally conserved at the Metazoa level, but the surrounding residues in the IDR are less conserved, increasing the relative score of the motif residues.

**Figure 4.**
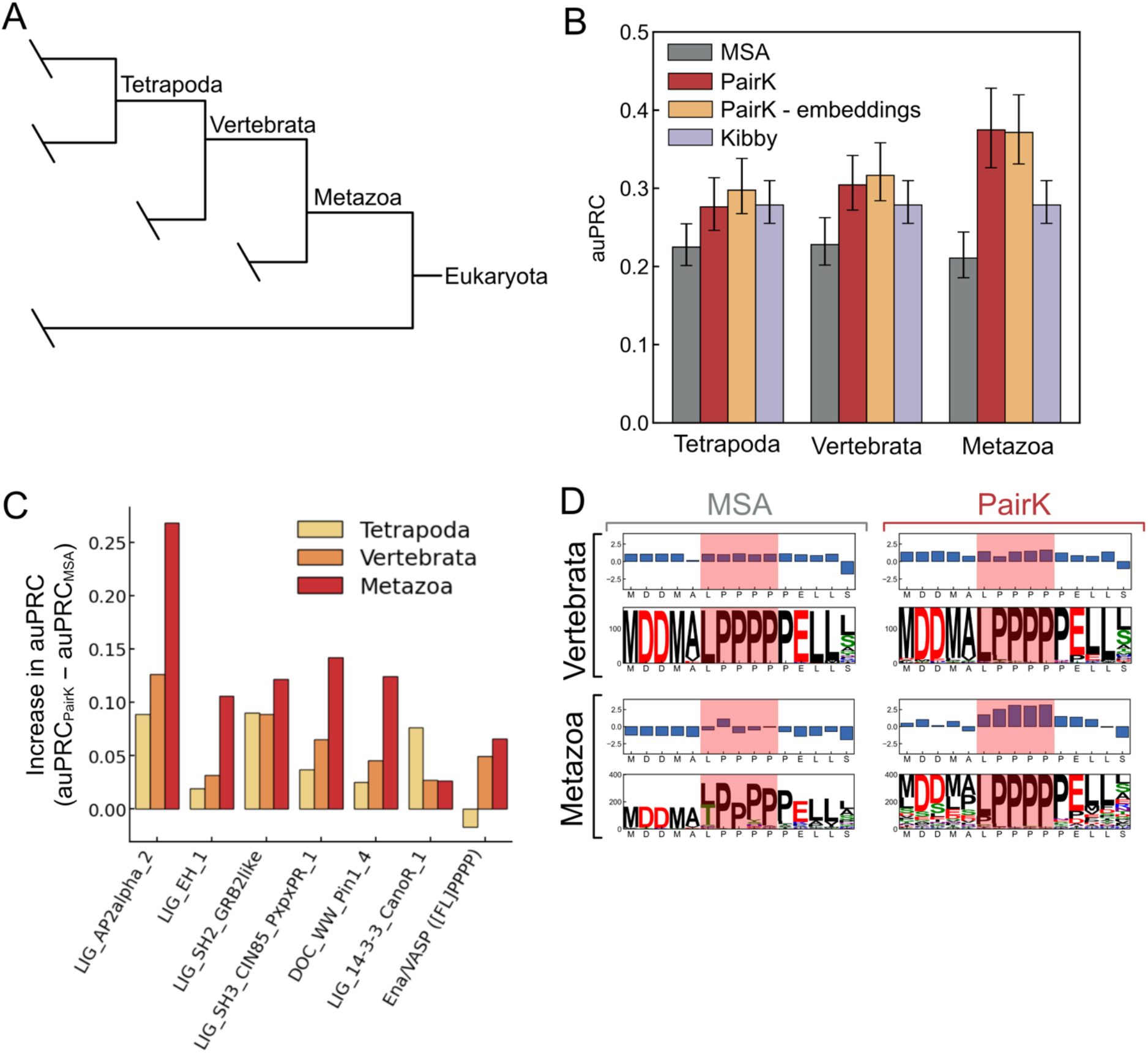
PairK better distinguishes real motifs from background matches for more divergent homologs. (A) Phylogenetic tree of Eukaryotes. (B) Performance (reported as auPRC) for the MSA and PairK methods at different phylogenetic levels. The performance of the Kibby method is replotted at each level for comparison with the other methods, however it is independent of phylogenetic level and only the query sequence is used in its calculation. Error bars are 95% confidence intervals from a bootstrap analysis. (C) The difference in auPRC score for PairK vs. the MSA method for individual motifs at different phylogenetic levels. (D) Example sequence logos and conservation scores for an experimentally verified SLiM from lamellipoden (RAPH1) that binds to the Ena/VASP EVH1 domain. The motif region is highlighted in red. For Vertebrata, the MSA and pairwise k-mer methods (*top*) perform similarly and show similar sequence profiles. For Metazoa, the MSA (*bottom left*) has a high fraction of gaps, indicated by white space in the logo, while the pairwise k-mer method (*bottom right*) indicates that the motif is still conserved in metazoans.

The enhanced accuracy of PairK, corroborated by benchmark results, opens avenues to uncover potentially significant but as-yet unvalidated SLiMs and to reveal conserved sequence attributes that contribute to SLiM binding. We scrutinized motif matches in the benchmark that were highly rated by PairK but poorly scored by the MSA method (detailed scores available in Supplementary Table S4) and identified several candidates for further study (illustrated in Figure 5).

**Figure 5.**
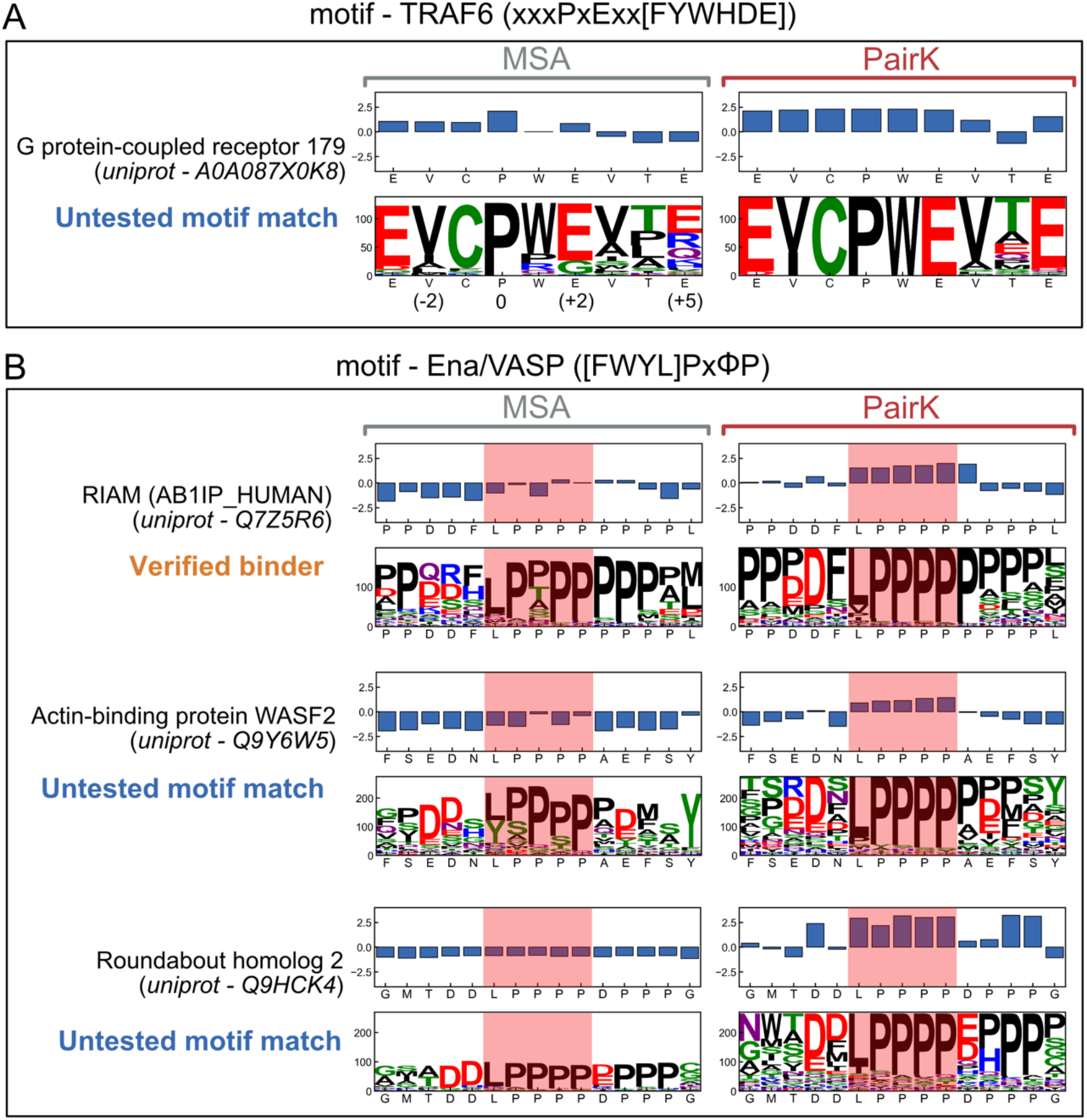
Sequence logos and conservation scores for examples from the benchmark. (A) TRAF6 motif match in G protein-coupled receptor 179 with homologs from the vertebrate level. (B) Motif matches for the Ena/VASP EVH1 domain (vertebrate level for RIAM, and metazoan level for WASF2 and Roundabout homolog 2). White space in the sequence logos indicates gaps in the alignment. The x-axis labels are the residues of the human sequence, which was the query sequence. For the MSA plots, the positions corresponding to the human residues were extracted from the MSA (as in Figure 1B) for easier visualization. For PairK plots, the sequence shown on the x-axis labels is the full query k-mer. In (B) a larger value of *k* was used (15) for the PairK method to show sequence flanking the motif. The motif residues are highlighted in red.

One example is sequence EVCPWEVTE from G protein-coupled receptor 179, which matches the TRAF6 preferred motif xxxPxExx[FYWHDE] (Figure 5A). TRAF6 is an E3 ubiquitin ligase that binds to cell-surface receptors and is involved in cellular processes including immunity and NF-kB signaling. Previous work suggests that TRAF6 highly disfavors binding to sequences with a proline at positions following the motif-required proline (position 0), because ligands form a beta sheet in complex with the domain (*45*– *47*). Conversely, TRAF6 favors binding to sequences with a negatively charged residue at the last position in the motif (position +5), based on the enrichment of this residue in known binding sequences (*46–52*). The MSA-derived sequence logo for a TRAF6 motif match in G protein-coupled receptor 179 indicates that features important for binding are not conserved in this protein. Glycine is observed at position +2, contrary to glutamate as mandated by the motif definition. Arginine and glutamine are also observed in position +5 instead of the preferred glutamate and aspartate. The presence of prolines in position +4 is incompatible with beta-sheet binding geometry. In contrast, when analyzed using PairK, G protein-coupled receptor 179 looks much more likely to bind TRAF6, as it is apparent that there are stretches of continuous sequence in all homologs that match the motif, and nearly all of those segments have glutamate at position +5 and no prolines after position 0. Little is known about the G protein-coupled receptor 179, so it is difficult to further assess whether it is a likely TRAF6 binding partner.

We identified two untested motif matches that are likely to be genuine Ena/VASP EVH1 binding partners: WASF2 and Roundabout Homolog 2 (ROBO2) (Figure 5B). Both proteins are biologically plausible Ena/VASP interaction partners, due to their roles in cytoskeletal processes. Moreover, ROBO2 is a paralog of the confirmed Ena/VASP binder ROBO1 (*53*), while WASF2 is a component of the WAVE complex, which plays a crucial role in lamellipodia formation (*54–56*). These motif matches are conserved according to PairK, but not when analyzed by MSA.

PairK can potentially unveil conserved sequence features that enhance binding affinity and specificity. The Ena/VASP motif from RIAM is a good example (Figure 5B). The MSA motif logo reveals a low conservation score for the first motif position and includes positively charged residues in the N-terminal flanking sequence. However, previous studies have shown that Ena/VASP EVH1 prefers negatively charged residues adjacent to the motif (*2, 57*). Furthermore, the first position of the motif is crucial for binding and engages a specific pocket on the EVH1 domain. In contrast to the MSA analysis, PairK reveals the conservation of negatively charged residues in the N-terminal flank and higher conservation of leucine at the first motif position.

## DISCUSSION

It is well known that disordered regions of proteins are challenging to align. Because MSAs form the basis for conservation scores, the quality of these alignments directly influences the reliability of conservation assessments, as illustrated in Figure 1. To detect conservation more reliably, we developed a multiple-sequence-alignment-free method termed PairK (pairwise k-mer alignment) (Figure 2) to quantify the conservation of short sequence fragments within IDRs (including SLiMs). We established a benchmark to evaluate the ability of our method to distinguish genuine SLiMs from background motif matches and showed that PairK significantly outperforms MSAs on this test.

PairK offers several advantages over typical MSA-based conservation methods when assessing SLiM conservation. One is its gapless nature. Deciding how much to penalize gaps in an MSA can be difficult and often arbitrary. PairK circumvents this issue by not allowing gaps at all. If there is no well-matching fragment in a homolog sequence, the scores are penalized through the mismatching residues rather than gap penalties. The fact that the pseudo MSAs are gapless greatly simplifies the interpretation of conservation at specific positions in the motif (Figure 5) and reduces artifacts from spuriously aligning high-scoring residues such as proline (Figure 1). It is more consistent with what is known about the binding of SLiMs to protein domains, which typically can’t accommodate gaps or insertions in the core motif because of the requirement to engage specific pockets on the binding domain.

One application for our tool is evaluating candidate SLiMs discovered in proteomic searches or experimental screens (*9, 58, 59*). Candidate motifs or binders from screens are not all biologically relevant. Thus, researchers are tasked with prioritizing sequences likely to yield significant biological insights for study, and sequence conservation emerges as a crucial factor in these decisions. MSA-based conservation scores can either underestimate or overestimate sequence conservation due to the quality of the underlying MSA. In contrast, PairK is less affected by artifacts resulting from aligning disordered regions (Figure 1) and is superior for distinguishing biologically relevant SLiMs from background matches (Figure 3). The protein language model-based conservation scoring method (the Kibby method (*24*)) also outperforms MSAs in our benchmark and has the advantage of not requiring homologous sequences at all. But Kibby did not perform as well as PairK for most motifs (Figure 3).

Another application of SLiM conservation scoring is generating hypotheses about important SLiM binding determinants, based on residue-level conservation. The identity and frequency of residues in homologs at each position within and around the motif (e.g., sequence logos as shown in Figure 5) are useful for this purpose. PairK is well-suited for this kind of analysis because it only utilizes gapless, continuous sequence fragments, and the pseudo MSAs can be analyzed or used to generate sequence logos that summarize motif features. This is not true of the Kibby method, which doesn’t provide the identity or frequency of residues in homologs at each position but only predicts a conservation score for each residue in a specific sequence.

We tested our conservation scoring methods on homologs from different phylogenetic levels (Figure 4). Notably, the performance of PairK improves with a broader phylogenetic range. As homologs diverge, the MSAs indicate that SLiMs are less conserved across metazoans. However, the enhanced performance of PairK implies that this apparent lack of conservation is due to declining MSA quality rather than reduced SLiM conservation. Alignments using Mafft, Muscle, and Clustalo at diverse phylogenetic levels (Supplementary Figure S2) support this observation, as the consistency of MSAs built using different tools diminishes with increased global divergence. Thus, our findings reveal an advantage of PairK—it enables the examination of SLiM conservation across broader phylogenetic ranges. This breadth enhances the conservation signal due to increased background divergence.

One consideration when using PairK is the choice of *k*. Adding residues flanking a potential motif (increasing *k*) could provide insight into the conservation of residues surrounding the motif. This could reveal residues outside of the core motif that are important for binding. Additionally, if a protein contains multiple motifs and the conservation of just one of them is of interest, the addition of flanking residues can help find the most similar motif in each homolog sequence. However, if *k* is too large, potentially irrelevant residues could influence the k-mer alignment and lower the overall quality of the results. The gapless nature of PairK assumes that residue positions are conserved due to contacts with the SLiM-binding domain. If *k* is too large and this assumption is broken, the resulting alignment quality will decrease. Our benchmark results suggest that PairK will perform differently for different SLiMs (Figure 3D). When first using our method, we advise running PairK on known instances of the SLiM of interest and comparing the conservation results with background matches to the motif, using a few different values for *k*. This will provide a sense of how conserved the specific SLiM instances are, how well the method separates true positives from background, and how much flanking sequence can be included without decreasing performance.

Several variations of our pairwise k-mer alignment method could be beneficial for specific analyses. A potential enhancement to our method is incorporating position weighting during pairwise alignment. In this approach, each query k-mer – homolog k-mer match score could be weighted based on individual positions, allowing users to include known SLiM information when selecting optimal scoring fragments from each homolog. For example, for the TRAF6 motif, xxxPxExx[FYWHDE] with *k*=9, scores from the “x” positions could be down-weighted in the alignment to prioritize selecting homolog fragments that match the essential P/E positions and the last position. Alternative scoring methods could easily be applied to the pseudo MSAs produced by pairwise k-mer alignment, for example, the regular expression-based score from (*18*). Additionally, PairK can be used without any prior knowledge of motifs or regular expressions, and thus could be used as an agnostic motif discovery tool.

We offer PairK as a free and publicly available Python package, available here: https://github.com/jacksonh1/pairk.

## METHODS

### General tools

We used Biopython (*60*) to facilitate sequence processing. Sequence logos were generated using the logomaker python package (*61*). Many of the plots were generated using the Seaborn (*62*) or matplotlib (*63*) python packages. The multiple sequence alignment image in Figure 1A was generated using Jalview (*64*).

### Pipeline for gathering and processing homologous sequences for conservation analysis

The following pipeline was used to generate all groups of homologous sequences in this study. For a protein of interest, (here called the query protein), we gathered precomputed orthologous groups at the specified phylogenetic level from a locally downloaded copy of the OrthoDB data v 11 (*36*). Any sequences that were shorter than 0.5 times the query sequence length were removed.

We further removed any sequences with non-amino acid characters (“X”, “x”, “*”, “J”, “B”, or “Z”). Next, we used the alfpy python package (v 1.0.6) (*22*) to calculate the sequence distance between the query sequence and all other sequences in the orthologous group (using google distance (*65*) between frequency vectors with a word size of 2). The sequence distances were then used to remove all but the closest sequence to the query sequence for each organism in the orthologous group (least divergent homolog), such that there remained one sequence for each organism. The remaining homologs were then clustered to 90% identity using CD-HIT (v 4.8.1) (*66, 67*) with the -g parameter. We redefined the representative sequence for the cluster containing the query protein to make sure that it was the cluster representative. The homolog sequences were then reduced to just the representative of each cluster. The final homolog sequences were then aligned with MAFFT (v 7.52) (*39*) unless otherwise specified. When Clustal Omega (v 1.2.3) (*41*) was used, we used default parameters. For Muscle (v 5.2) (*40*), default parameters were used except we added the -super5 flag. The pipeline used to generate the homolog groups is available at https://github.com/jacksonh1/slim_conservation_orthogroup_generation.

### Definition of disordered regions

To define the disordered regions in a query protein, we used IUPred2A (*11, 12*) to calculate disorder scores for the sequence. We defined the IDRs as regions in the query sequence where the IUPRED scores were above 0.4. If two IDRs were separated by fewer than 11 residues, we merged them into one IDR. If an IDR was shorter than 8 residues, it was discarded.

### Column-wise conservation scores

To calculate conservation scores for MSA columns and PairK’s pseudo MSA columns, we used the python script from Capra et. al (*42*), which we trivially modified for compatibility with modern python tools. Unless otherwise specified, conservation scores were calculated for each column of an MSA (or pseudo MSA) using the property entropy score from the Capra et. al script. The residue groups (V, L, I, M), (F, W, Y), (S, T), (N, Q), (H, K, R), (D, E), (A, G) were treated as equivalent amino acids (*68*). For Figure 1 and S2, the Shannon entropy score from the Capra et. al script was used. For both the Shannon entropy and property entropy, a gap penalty was applied to the final score for the column (as in Capra et. al), where the score was multiplied by the fraction of the column that was gaps (here termed gap fraction). The scores were inverted and normalized such that the values ranged from 0 to 1, with 1 being maximally conserved and 0 reflecting no conservation. Both Shannon entropy and property entropy showed similar performance on the benchmark (Supplementary Figure S1D)

### Variability in MSAs generated by different methods for regions containing SLiMs (Figure 1C and Supplementary Figure S2)

The homolog group pipeline described above was used to gather homologous sequences for proteins containing verified SLiMs. The same set of SLiMs was used as was used in the benchmark (TP set, described below) except for LIG_14-3-3_CanoR_1, which was removed due to its variable-length regular expression. MSAs were produced with the final homolog sequences using MAFFT, Muscle, and Clustal Omega (see pipeline methods above for MSA details). From the MSAs, Shannon entropy conservation scores were calculated for the columns corresponding to the defined positions of the motif, *i*.*e*., any position in the regular expression not defined as “x” (x = any residue) (see Supplementary Table S1 for motif position masks used).

### MSA-based conservation score

To calculate conservation scores from an MSA, the property entropy score was first calculated for each column in the alignment (described above). For positions within the IDR (as defined by the query sequence, described above), we converted the scores to z-scores. The background score distribution used for the z-score calculation was every score in the IDR whose column had a gap fraction less than 0.2. Thus, the z-score for each position was the column score minus the mean of the background distribution divided by the standard deviation of the background distribution. The reported score for the motif was the average z-score of the defined motif residues (see Supplementary Table S1 for motif position masks used).

### Kibby conservation scores

The conservation score predictor, Kibby, from Yeung et. al. (*24*) generates per-residue conservation scores for input sequences. It does not require an alignment or homologous sequences. We used the Kibby conservation score predictor to generate conservation scores for the full-length query sequences in the benchmark datasets. The Yeung et al. script conservation_from_fasta.py was used with default parameters (language model esm2_t33_650M_UR50D) and ‘-device=cuda’. The resulting conservation scores were converted to z-scores in the same manner as the MSA-based conservation scores, using the conservation scores of the IDR residues as the background score distribution. No gap fraction mask was used as there were no gaps or alignment involved. The reported score for the motif was the average z-score of the defined motif residues (see Supplementary Table S1 for motif position masks used).

### Pairwise k-mer alignment (PairK) method

To run PairK for a query IDR, we had to obtain the corresponding IDR in each homolog. For a direct comparison with the MSA-based method, we extracted the homolog IDRs from an MSA by slicing the region of the alignment corresponding to query IDR. Thus, the same IDR sequences that are used in the MSA-based score calculations are used as input for PairK. The IDRs were then de-aligned by removing all the gaps. From the query IDR, overlapping k-mer sequences were generated using a sliding window approach. Starting from the first position in the sequence, the sequence within a window of length *k* was recorded as the k-mer for that position. This process was repeated for each position in the query IDR until the window reached the end of the sequence. For each k-mer in the query IDR (k-mer_q_), a gapless pairwise “pseudo MSA” was constructed using the following procedure. For each homolog IDR, k-mers were generated (k-mer_h_). The k-mer_h_ that best matched the query k-mer_q_ was then identified by calculating an alignment score for all possible k-mer_q_ - k- mer_h_ alignments (gapless) using a scoring matrix. For each residue match in the k-mer_q_ - k-mer_h_ alignment, the match score was retrieved from the scoring matrix, and the sum of the match scores was taken as the alignment score. The pseudo MSA for each k-mer_q_ was constructed by collecting the highest scoring k-mer_h_ from each homolog IDR. In instances where there was more than one best-scoring k-mer_h_, the first instance (most N-terminal) was selected (behavior of the numpy argmax function (*69*)). Unless otherwise specified, we used the EDSSMat50 substitution matrix (*44*), which was built for disordered regions, as the scoring matrix. We also tested the Blosum62 matrix (*70*) and Grantham distance matrix (*71*). The Grantham distance matrix values were normalized to the range 0 - 1 by subtracting from each matrix element the lowest value in the matrix and dividing by the difference between the highest and lowest matrix values. The normalized matrix was then converted to a similarity matrix by subtracting each matrix element from 1. All three matrices showed very similar performance, however the EDSSMat50 performed best (Supplementary Figure S1).

For the ESM-embedded version of PairK, we used a similar procedure as for the normal PairK method, except residue embeddings from ESM2 (*35*) were used to select the best matching homologous subsequences for each k-mer. In this version of the method, ESM2 residue embeddings were computed for the full-length query sequence and homolog sequences, using the esm2_t33_650M_UR50D model. The start and stop tokens were then removed leaving a tensor of dimension n x 1280, where n is the number of residues in the sequence. Thus, each amino acid in the sequence had an associated 1280-dimension feature vector. When the query IDR was split into k-mer_q_ segments, the corresponding residue embeddings were also sliced out of the query embedding tensor, *i*.*e*., each k-mer_q_ was associated with a tensor slice, T_ij_^q^ ∈ ℝ ^k x 1280^. For each homolog IDR, k-mer_h_ segments and their associated embedding slices were also generated, T_ij_^h^ ∈ ℝ ^k x 1280^.For each k-mer_q_ and for each homolog IDR, the best matching k-mer_h_ was determined by calculating the Euclidean distance between all possible T_ij_^q^ − T_ij_^h^ pairs. The pseudo MSA for each k-mer_q_ was constructed by collecting the k-mer_h_ from each homolog IDR with the lowest Euclidean distance to that k-mer_q_.

After constructing pseudo MSAs for all query IDR k-mers (by either version of PairK) we calculated conservation scores for every column in every pseudo MSA (see “column-wise conservation scores” above). The conservation scores were then converted to z-scores, using all columns in all of the pseudo MSAs as the background distribution. The reported score for the motif was the average z-score of the defined motif residues (see Supplementary Table S1 for motif position masks used). We have made PairK available as a python tool (https://github.com/jacksonh1/pairk).

### SLiM conservation benchmark

The code used to preprocess the data sources and generate the benchmark is available at https://github.com/jacksonh1/slim_conservation_benchmark.

### Preprocessing of ELM data

We downloaded the following data tables from the ELM (downloaded on February 9, 2024) (*38*): the SLiM classes table that contains the regular expression for each motif class and the SLiM instances table, which contains the experimentally verified SLiM instances. The fasta file containing the full-length sequences for each instance was also downloaded to obtain the amino-acid sequence of the SLiM from the positions provided in the instance table. We then removed the small number of instances for which the SLiM sequence did not match the regular expression provided for the SLiM. The sequences used for the ELM annotations and the sequences used in OrthoDB can be slightly different, e.g., when the two databases used different sequence sources or different versions of the same sequence. Therefore, we had to map the ELM instance annotations to the corresponding proteins in OrthoDB. To do so, we mapped the UniProt ID for each instance in the ELM to its corresponding protein in OrthoDB using the homolog group pipeline tools described above and removed the small fraction of instances that could not be mapped. Finally, we found the SLiM positions within the OrthoDB sequence by searching for the ELM SLiM sequence within the full-length OrthoDB sequence. To ensure that only one copy of the SLiM was found in the OrthoDB sequence, we included 15 residues flanking each side of the ELM SLiM sequence in the search. No sequences had multiple copies of the sequence used in the search. The small number of entries that did not contain a perfect match to the search sequence were discarded.

### Benchmark - true positive set

We chose the following SLiMs from the ELM to use in the benchmark: DOC_WW_Pin1_4, LIG_AP2alpha_2, LIG_EH_1, LIG_SH2_GRB2like, LIG_14-3-3_CanoR_1, and LIG_SH3_CIN85_PxpxPR_1, based on the number of verified instances that were available. To generate the true positive set, we first filtered the preprocessed ELM data (described above) to include only SLiM instances annotated as true positives (most of the instances in the table). Any instances not annotated as *Homo sapiens, Mus musculus*, or *Rattus norvegicus* were then removed. Because we used only human sequences to search for background motif matches, we only included these closely related organisms in the true positive (TP) set. For each protein, we determined the disordered regions as described above. Any instances not in an IDR were removed. We added Ena/VASP EVH1 domain and the TRAF6 MATH domain to the benchmark but manually curated the verified SLiM instances for these domains from the literature (Supplementary Tables S2 - S3) because we noticed either missing or incorrect annotations in the ELM for those SLiMs. The instances for Ena/VASP and TRAF6 were filtered in the same manner as the ELM instances. TRAF6 was removed from the overall analysis of the benchmark, due to the small number of true positives, but we calculated conservation scores for the motif anyway to include in Supplementary Table S2.

### Benchmark – background set

To generate the background matches (BG set), we retrieved all human sequences in OrthoDB. Although OrthoDB purports to remove duplicate and isoform sequences in each organism, we found some of these. Therefore, we clustered the set of all human sequences in OrthoDB to 95% identity using CD-HIT (using the arguments -c 0.95 and -g 1) and kept only the representative sequence from each cluster. Clusters containing a protein with a true positive SLiM were removed. We used the remaining set of human sequences to search for background matches (∼19,000 sequences total). To find the background motif matches, we first defined the disordered regions in the background sequences as described above. For each SLiM in the TP set, the motif regular expression was used to search the background sequences. Sequences matching the regular expression and in a disordered region were added to the background set for that SLiM. Finally, the full set of ELM instances (preprocessed) was used to remove any motif match in the BG set that overlapped with any verified SLiM instance. In cases where the BG set was very large for a given motif, the BG set was randomly subsampled to reduce computation time. Finally, sequences longer than 5000 residues were removed from all sets. Final counts of the TP/BG sets for each motif are provided in Supplementary Table S1.

### Benchmark scoring

To evaluate the performance of the MSA and PairK method on the benchmark, conservation scores were calculated for all motif matches in the benchmark for homolog groups collected at the Tetrapoda, Vertebrata, and Metazoa phylogenetic levels (constructed using the pipeline described above). For MSA conservation scores, the motif match (either BG or TP) was discarded if there were fewer than 20 scored columns in the background score distribution used to calculate the z-scores, *e*.*g*., if too many columns exceed the gap fraction cutoff value of 0.2 or for very short IDRs with fewer than 20 columns. For PairK, the motif match was discarded if there were fewer than 50 points in the background score distribution used to calculate the z-scores or fewer than 10 unique k-mers in the query IDR. The final reported conservation score for the motif for both methods is the average z-score of the defined motif residues (positions that are not ‘x’ in the motif). To be included in the final benchmark, we required each motif match to have a valid score for all methods at all of the phylogenetic levels. For example, if a motif match failed the MSA score method at the Metazoa level due to having too few background scores for a z-score calculation, it was removed entirely from the benchmark even if it passed at the other levels. The full benchmark table with conservation scores is provided in Supplementary Table S2.

### Benchmark bootstrapping analysis

To test the robustness of the auPRC scores from the benchmark, we performed a bootstrapping analysis. We generated 1000 bootstrap replicates of the benchmark scores by random sampling with replacement. In each replicate, the number of scores for each SLiM and the number of true positives and background matches for each SLiM were kept the same as for the original benchmark scores. We calculated auPRC scores for each bootstrap replicate for the entire benchmark (Figure 4) and for each SLiM (Figure 3). We then calculated 95% confidence intervals from the bootstrap replicate auPRCs using the percentile function from the numpy python package (*69*).

## ACKNOWLEDGEMENTS

This work was supported by the National Institutes of Health under Award Number R35GM149227 and F32GM137510. The content is solely the responsibility of the authors and does not necessarily represent the official views of the National Institutes of Health. We thank Foster Birnbaum for help accelerating the embedding distance functions.

## Notes

### Competing Interest Statement

The authors have declared no competing interest.

