## Supplementary Information for "PairK: Pairwise k-mer alignment for quantifying protein motif conservation in disordered regions"

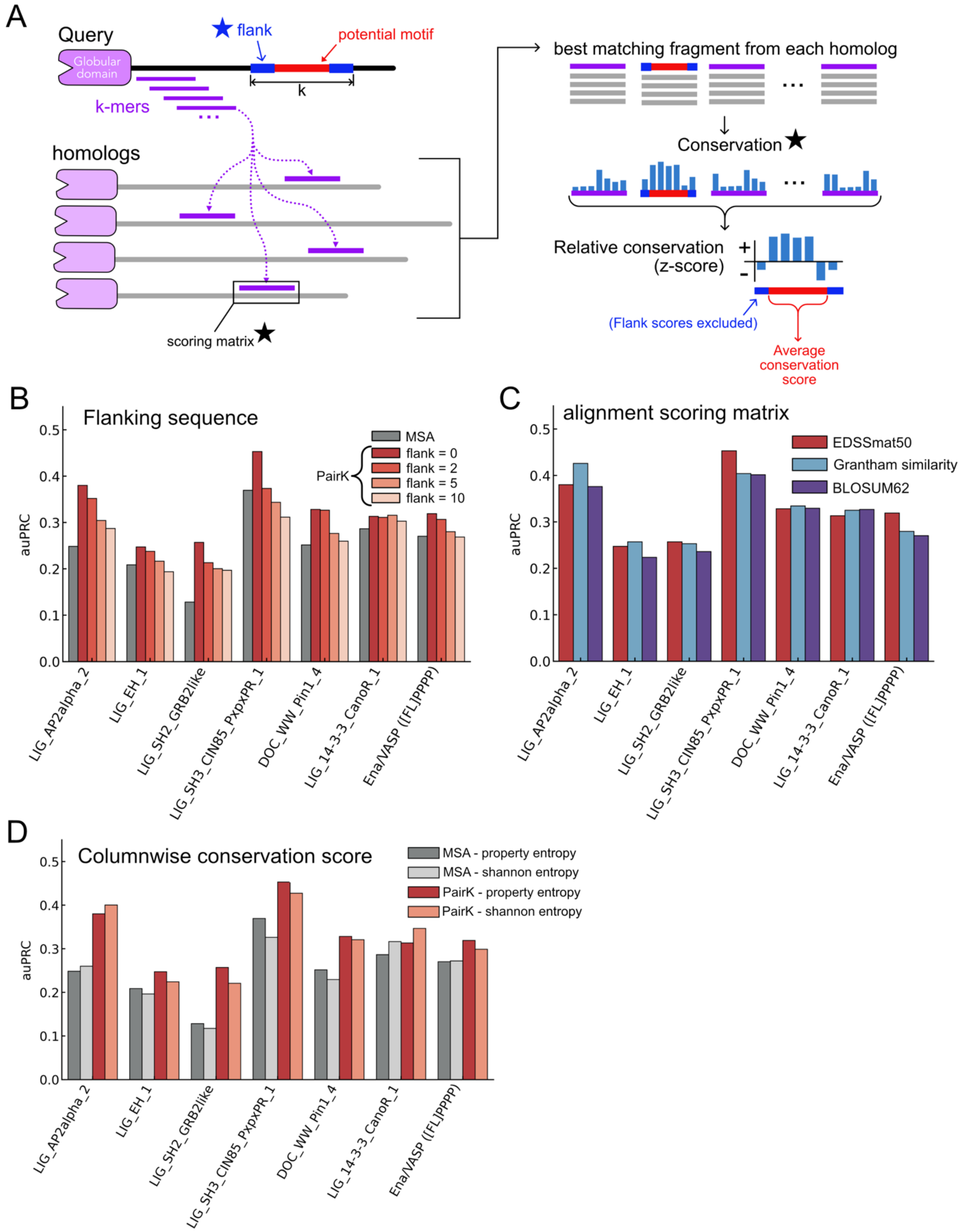


**Figure S1.** Effect of flanking sequence, scoring matrix, and conservation score method on PairK performance. (A) The PairK method. Stars indicate where the method is changed in parts B-D. The performance of the method is shown when adding residues flanking the potential motif for the alignment step (i.e. increasing k) (B), changing the alignment scoring matrix (C), and using different column-wise conservation scoring methods (D). Column-wise scoring methods used: property entropy and Shannon entropy (*1*). Scoring matrices: Blosum62 (*2*), EDSSmat50 (*3*), and the Grantham matrix converted to a similarity matrix (*4*). Homolog sequences were gathered at the Vertebrata level.

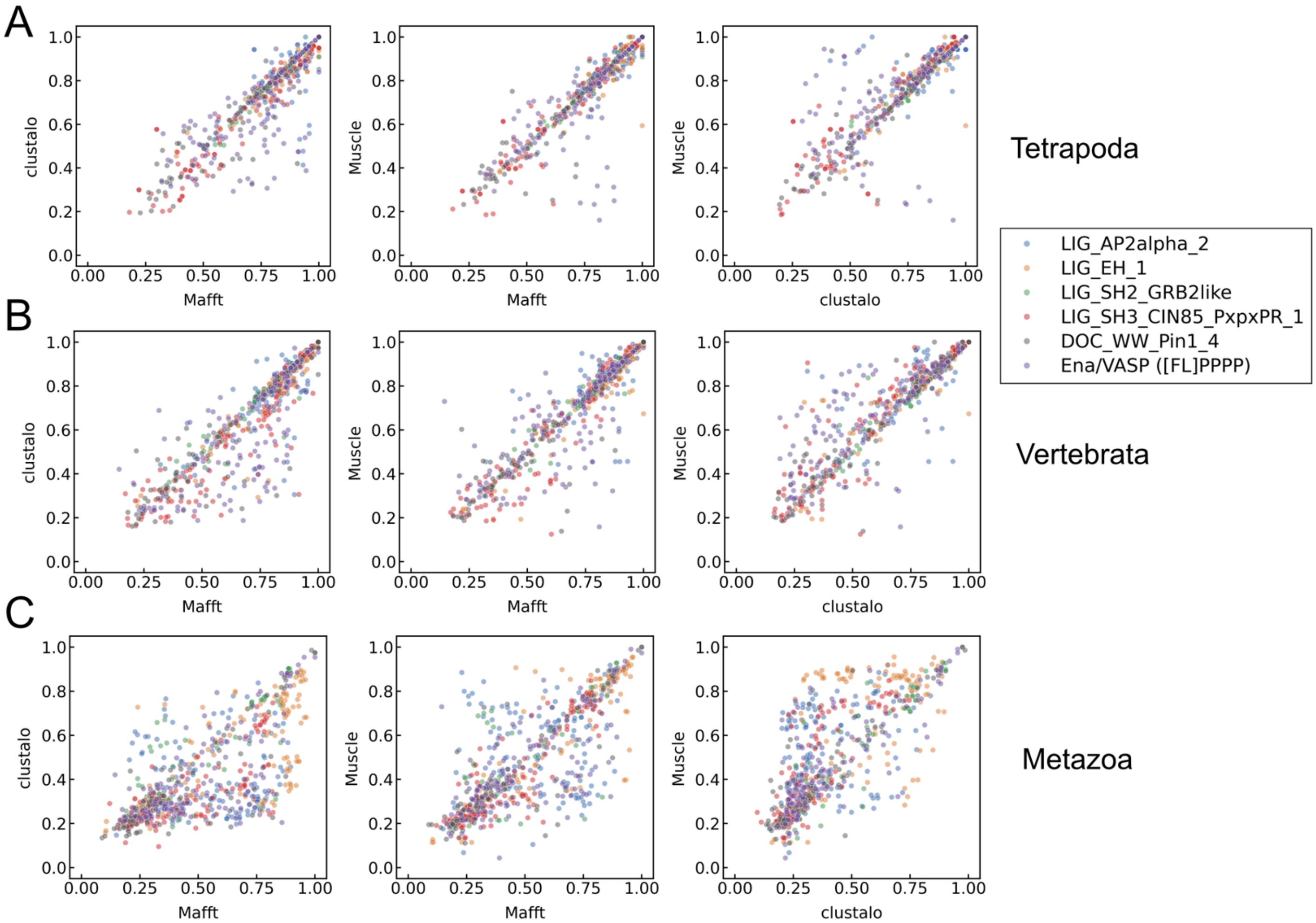


**Figure S2.** The correlation between conservation scores of residues in experimentally verified SLiMs from MSAs produced by different alignment algorithms. The underlying homolog sequences were collected at the Tetrapod (A), Vertebrate (B), or Metazoa (C) level. Data are from 236 verified SLiM instances (721 residues).

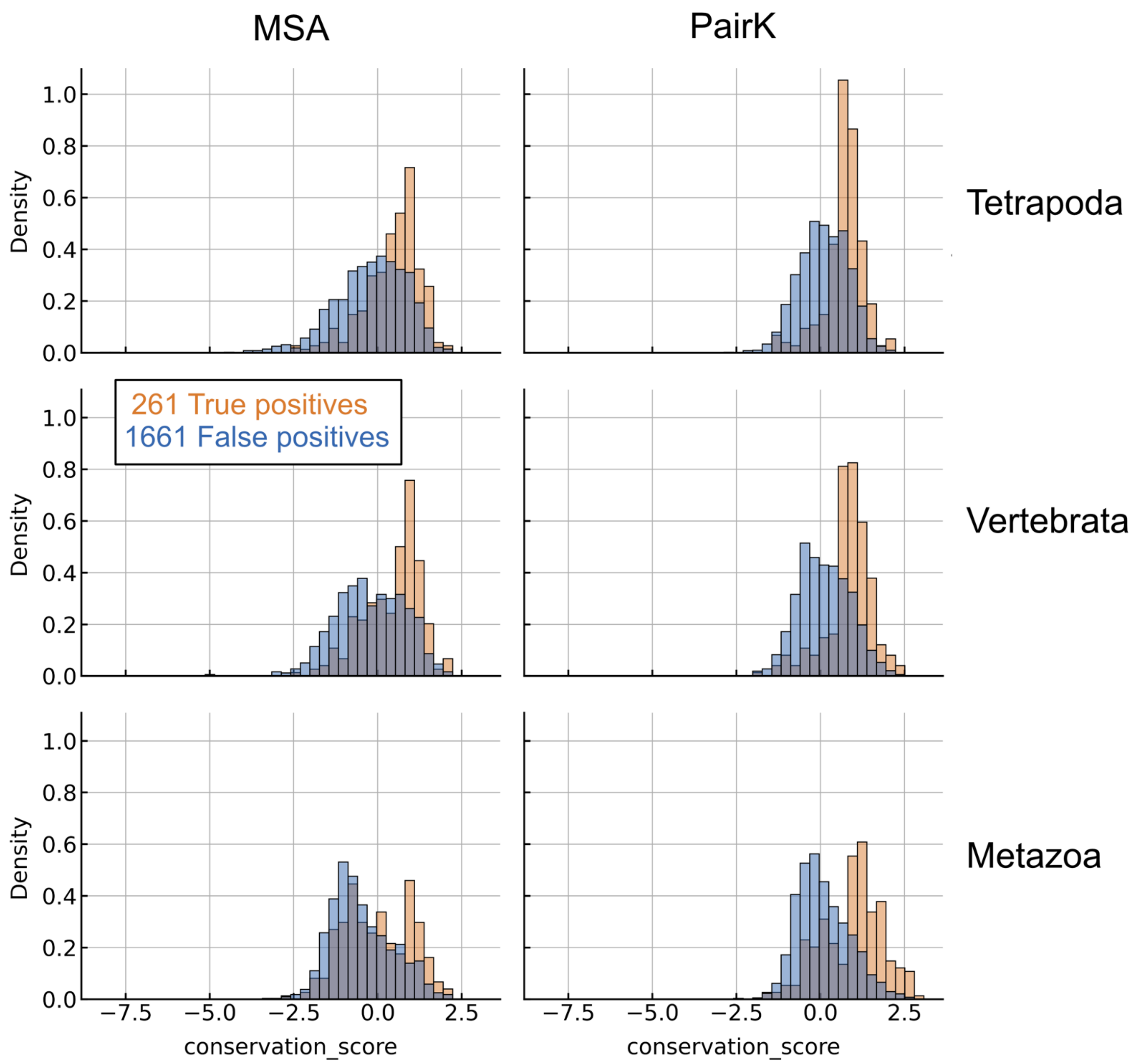


**Figure S3**. Benchmark conservation score distributions (density) from homolog groups retrieved at the Tetrapoda, Vertebrata and Metazoa levels. True positives are shown in orange and background matches are shown in blue.

### Supplementary Tables

**Table S1.** Counts of SLiM instances in the benchmark for each motif.

| SLiM | background | true positives | motif regular expression | mask‡ |
|---|---|---|---|---|
| DOC_WW_Pin1_4 | 242 | 42 | ...([ST])P. | [0 0 0 1 1 0] |
| Ena/VASP ([FL]PPPP) | 241 | 34 | [FL]PPPP | [1 1 0 1 1] |
| LIG_14-3-3_CanoR_1 | 246 | 41 | R[^DE]{0,2}[^DEPG]([ST])(([FWYLMV].)\|([^PRIKGN]P)\|([^PRIKGN].{2,4}[VILMFWYP])) | No mask |
| LIG_AP2alpha_2 | 241 | 49 | DP[FW] | [1 1 1] |
| LIG_EH_1 | 269 | 49 | .NPF. | [0 1 1 1 0] |
| LIG_SH2_GRB2like | 210 | 18 | (Y)([EDST]\|[MLIVAFYHQW])N. | [1 1 1 0] |
| LIG_SH3_CIN85_PxpxPR_1 | 212 | 28 | P.[AP].PR | [1 0 1 0 1 1] |
| TRAF6* | 199 | 7 | ...P.E..[FYWDE] | [0 0 0 1 0 1 0 0 1] |

*TRAF6 was removed from global analyses (*e.g.* auPRC calculations) due to the low number of true positive instances.
‡Positions in the mask array with a 0 were not included in the average conservation score of the motif, whereas positions with a 1 were included. For example, for the sequence DPW and a mask of [1 0 1], the conservation score of the motif would be reported as the average of the first (D) and last (W) residue scores.

**Table S2**. Manually curated verified Ena/VASP EVH1 binding partners (*5–20*). Interaction data are at the protein level so, for each protein, any FPPPP and LPPPP sequence within an IDR was considered a true positive.

| Name | Uniprot ID | reference DOI | OrthoDB id |
|---|---|---|---|
| AB1IP_HUMAN | Q7Z5R6 | 10.1016/j.devcel.2004.07.021 | 9606_0:00294e |
| ABI3_HUMAN | Q9P2A4 | 10.1016/j.devcel.2014.08.001 | 9606_0:003dae |
| ANK3_HUMAN | Q12955 | ELM - 10.1093/nar/gkad1058 | 9606_0:0027f1 |
| FAT1_HUMAN | Q14517 | 10.1038/sj.emboj.7600380 | 9606_0:00122c |
| FBLI1_HUMAN | Q8WUP2 | 10.1074/jbc.M512107200 | 9606_0:000661 |
| FYB1_HUMAN | O15117 | 10.1083/jcb.149.1.181 | 9606_0:0015fb |
| LPP_HUMAN | Q93052 | 10.1091/mbc.11.1.117 | 9606_0:000d90 |
| NHSL1_HUMAN | Q5SYE7 | 10.7554/eLife.70680 | 9606_0:001b40 |
| PALLD_HUMAN | Q8WX93 | 10.1002/cm.10173 | 9606_0:00141f |
| PCARE_HUMAN | A6NGG8 | 10.7554/eLife.70680; 10.1073/pnas.1903125117; 10.1038/ncomms11491 | 9606_0:00094d |
| RAPH1_HUMAN | Q70E73 | 10.1016/j.devcel.2004.07.024 | 9606_0:000b76 |
| ROBO1_HUMAN | Q9Y6N7 | ELM - 10.1093/nar/gkad1058; 10.1016/s0092-8674(00)80883-1 | 9606_0:000e6f |
| SHIP2_HUMAN | O15357 | 10.1083/jcb.201501003 | 9606_0:002a4b |
| SHRM3_HUMAN | Q8TF72 | 10.7554/eLife.70680; 10.1242/dev.045369 | 9606_0:0012c3 |
| VINC_HUMAN | P18206 | ELM - 10.1093/nar/gkad1058 | 9606_0:002935 |
| XIRP1_HUMAN | Q702N8 | ELM - 10.1093/nar/gkad1058; 10.1016/j.yexcr.2006.03.015 | 9606_0:000f47 |
| ZYX_HUMAN | Q15942 | ELM - 10.1093/nar/gkad1058; 10.1074/jbc.M001698200 | 9606_0:001d2c |

**Table S3**. Manually curated verified TRAF6 MATH domain interactions (*21–27*).

| Name | Uniprot ID | SLiM sequence | reference DOI | OrthoDB id |
|---|---|---|---|---|
| CD40 | P25942 | KQEPQEINF | 10.1073/pnas.96.4.1234, 10.1074/jbc.274.20.14246 | 9606_0:004882 |
| TIFA | Q96CG3 | SSSPTEMDE | 10.1002/cbic.201800436 | 9606_0:001440 |
| MAVS | Q7Z434 | CHGPEENEY | 10.1074/jbc.M115.666578 | 9606_0:00486f |
| TICAM1 | Q8IUC6 | CQEPEEMSW | 10.1073/pnas.0308496101, 10.4049/jimmunol.171.8.4304 | 9606_0:004368 |
| IRAK2 | O43187 | SNTPEETDD | 10.1038/nature00888 | 9606_0:000e31 |
| IRAK1 | P51617 | PPSPQENSY | 10.1038/nature00888 | 9606_0:004fa3 |
| IRAK1 | P51617 | PNQPVESDE | 10.1038/nature00888 | 9606_0:004fa3 |
| IRAK1 | P51617 | RQGPEESDE | 10.1038/nature00888 | 9606_0:004fa3 |
| IRAK3 (IRAK-M) | Q9Y616 | PSIPVEDDE | 10.1038/nature00888 | 9606_0:0031e9 |
| mouse TNFRSF11A (RANK) | O35305 | RKIPTEDEY | 10.1038/nature00888 | 10090_0:000361 |
| mouse TNFRSF11A (RANK) | O35305 | FQEPLEVGE | 10.1038/nature00888 | 10090_0:000361 |
| mouse TNFRSF11A (RANK) | O35305 | GNTPGEDHE | 10.1038/nature00888 | 10090_0:000361 |
